# Comparative genomics and transcriptomics on salt tolerance of *Vigna luteola*

**DOI:** 10.1101/2025.05.21.653682

**Authors:** Yurie Iki, Fanmiao Wang, Kosuke Ito, Takanori Wakatake, Keitaro Tanoi, Ken Naito

## Abstract

*Vigna luteola*, a wild legume species, shows remarkable variation in salinity tolerance across its natural habitats, with coastal populations exhibiting high tolerance and riverbank populations being sensitive. This intraspecific variation provides a valuable system for investigating the genetic basis of salt tolerance. A major QTL for salt tolerance was previously identified by crossing salt-tolerant and salt-sensitive accessions, but the responsible genes remain unknown. In this study, grafting experiments between the two accessions revealed that the root plays a primary role in salt tolerance by suppressing Na⁺ transport to the shoot. We then conducted root transcriptome analysis and identified four candidate genes located within the QTL and highly expressed under salt stress in the tolerant accession: *CBL-INTERACTING PROTEIN KINASE 6 (CIPK6)*, *CAFFEOYL SHIKIMATE ESTERASE (CSE)*, *FCS-LIKE ZINC FINGER PROTEIN 13 (FLZ13)*, and *DROUGHT-INDUCED 21 (DI21)*. Promoter analysis revealed that the *CIPK6* promoter contains transcription factor binding motifs unique to the salt-tolerant accession, which may contribute to its high expression under salt stress. These findings suggest that *CIPK6* is regulated by *cis*-regulatory differences and is the most promising candidate for the salt-tolerance QTL. The identified genes in this study provide a foundation for developing salt-tolerant crops in the future.

## Introduction

Groundwater depletion will be a serious problem in world agriculture and significantly impact nations that rely on imported products for their staple food supply (Wada *et al*. 2010; Dalin *et al*. 2017). Thus, it is urgent to develop salt-tolerant crops to realize agriculture with saline water.

Although hundreds of genes potentially involved in salt tolerance have been identified, it remains unclear how to utilize them effectively. Since manipulation of all these genes is impractical, identifying the best combination is essential. Furthermore, it is also critical to determine the specific organs or tissues where these genes should be expressed, as well as the optimal timing for their expression. It would be almost impossible to test all these possibilities in model plants.

Instead, we may find a direct answer in wild species that are adapted to a saline environment. As one such species, we have focused on *Vigna luteola* (Jacq.) Benth. It mainly lives in tropical marine beaches in West Africa and the Americas, and some accessions can survive a condition of 400 mM NaCl for more than 4 weeks (Yoshida *et al*. 2020). Interestingly, some populations of this species have settled in riverbanks and are not salt tolerant (Yoshida *et al*. 2020; Noda *et al*. 2025). We have previously crossed the salt-tolerant and salt-sensitive accessions and identified a single major QTL involved in salt tolerance (Chankaew *et al*. 2014). Other previous studies revealed that the salt-tolerant accession allocates more Na^+^ in the root (Noda *et al*. 2022; Wang *et al*. 2024), whereas the salt-sensitive one allocates more Na^+^ in the shoot (Wang *et al*. 2024; Noda *et al*. 2025). In addition, the salt-tolerant accession consistently allocates Na^+^ to the topmost fully expanded leaves, suggesting the “taking turns” system of Na-loading leaves (Noda *et al*. 2022). These results indicate that the salt-tolerant accession possesses distinct salt-tolerant mechanisms in the root and shoot.

However, it remains unclear whether the root contributes more to salt tolerance or the shoot does. Moreover, the genes responsible for salt tolerance in *V. luteola* have not yet been identified.

In this study, we first conducted grafting experiments between the salt-tolerant and salt-sensitive accessions and examined their salt tolerance and Na^+^/K^+^ allocation. Our results suggest that the root plays a more critical role in salt tolerance and suppressing Na^+^ transport to the shoot under salt stress. Based on these findings, we analyzed the root transcriptome data (Noda *et al*. 2025) and identified four candidate genes that met the following three criteria: (1) located within the major QTL region related to salt tolerance, (2) highly expressed in the salt-tolerant accession under salt stress, and (3) homologous to genes with functions potentially contributing to salt tolerance. Promoter regions of the four candidate genes were then analyzed to identify differences in *cis*-regulatory motifs potentially involved in the different expression patterns between the two accessions. As a result, *CIPK6* was considered the most promising candidate gene for root-mediated salt tolerance mechanisms. The candidate genes identified in this study could be targets for developing salt-tolerant crops in the future.

## Materials and Methods

### Plant materials

Two accessions of *Vigna luteola* were used: a salt-tolerant accession, JP233389, which inhabits beaches (hereafter referred to as tolerant accession), and a salt-sensitive accession, JP235855, which inhabits riverbanks (hereafter referred to as the sensitive accession). These seeds were obtained from the Research Center of Genetic Resources, National Agriculture and Food Research Organization (NARO), Japan (NARO Genebank: https://www.gene.affrc.go.jp/index_en.php).

Plants were grown as described previously (Wang *et al*. 2024). Briefly, germinated seeds were transplanted into a hydroponic solution and grown in a growth chamber. The hydroponic solution was used at double the concentration compared to Wang et al. (2024) to minimize nutritional deficiency stress during the salt tolerance test and during ion concentration analysis. Specifically, we used OAT House No.1 and OAT House No.2 (Otsuka Chemical Co., Japan) at double the standard concentrations, resulting in final concentrations of 37.2 mEq L^−1^ N, 10.2 mEq L^−1^ P, 17.2 mEq L^−1^ K, 16.4 mEq L^−1^ Ca, and 6.0 mEq L^−1^ Mg.

### Grafting experiments

Germinated seedlings were transplanted to a hydroponic culture as described above. After 1–2 days, seedlings were cut in a V-shape (Supplemental Figure 1) and grafted using 12.5 mm wide Micropore surgical tape (3M Japan Limited, Tokyo, Japan). Four grafting combinations (scion/rootstock) were produced: tolerant/tolerant (T/T), tolerant/sensitive (T/S), sensitive/tolerant (S/T), and sensitive/sensitive (S/S). Shoots and roots were taken from different plants even if they were the same accessions (T/T or S/S). After grafting, plants were returned to the original hydroponic culture and were initially maintained in a high-humidity environment after grafting to prevent desiccation.

### Salt tolerance test of grafted plants

After the third leaf fully expanded, grafted plants were transferred to the hydroponic culture containing 200 mM NaCl. After 5, 8, 12, and 15 days of salt treatment, wilt scores were evaluated by three to four persons using a scale of 1, 3, 5, 7, and 9, corresponding to 0%, 1–25%, 25–50%, 50–75%, and 100% wilting, respectively.

### Analysis of Na^+^ and K^+^ allocation

The methods followed those described previously by Noda et al. (2022) with minor modifications. Briefly, germinated plants were transferred to the hydroponic culture containing double the standard concentrations of OAT House No.1 and OAT House No.2. After the first leaf fully expanded, tolerant and sensitive accessions were treated with 200 mM NaCl in the hydroponic culture for three days. Subsequently, shoots and roots were sampled, dried, and digested with 60% and 69% HNO3, respectively. Then, the sodium (Na) and potassium (K) contents were measured using inductively coupled plasma-mass spectrometry (ICP-MS, NexION 350S, PerkinElmer, Waltham, MA, USA).

Grafted plants were also subjected to the same experiments after the third leaf fully expanded. The graft sites, covered with surgical tape, were not sampled due to the difficulty of removing the tape.

### DNA extraction, genome sequence, and assembly of the sensitive accession

Genome sequencing and assembly for the tolerant accession were previously conducted by Noda et al. (2025). Therefore, in this study, we did genome sequencing and assembly only for the sensitive accession.

As described in (Naito 2023), high molecular weight DNA was extracted from unexpanded leaves with the NucleoBond HMW DNA kit (MACHEREY-NAGEL GmbH & Co. KG, Düren, Germany), size-selected with Short Read Eliminator XL kit (Pacific Biosciences of California, Inc., California, USA). Sequencing libraries for nanopore sequencing were prepared with SQK-LSK110 (Oxford Nanopore Technologies KK, Tokyo, Japan). Each library was then loaded to PromethION R9.4.1 Flowcell (Oxford Nanopore Technologies KK, Tokyo, Japan) and was run for 72 h.

The obtained raw data was transformed into fastq format with guppy-5.0.7. Short reads were also obtained with HiSeq 4000 (Illumina, San Diego, USA), which was provided as a customer service by GeneBay, Inc. (Yokohama, Japan).

Draft assembly was performed with necat-0.0.1 (Chen et al., 2021) with default parameters except “PREP_OUTPUT_COVERAGE=60” and “CNS_OUTPUT_COVERAGE=40”. Polishing was done with racon-1.4.3 (Vaser et al., 2017) and medaka-1.0.3 (https://github.com/nanoporetech/medaka) twice each. Further polishing was also done with hypo-1.0.3 using short reads. Scaffolds were generated using RagTag (Alonge *et al*. 2022) using the genome of the tolerant accession as a reference. Any controversies between the contigs and linkage maps were regarded as misassemblies and manually fixed as described by Sakai et al. (2015). Genome completeness of the assembly was evaluated using BUSCO (5.8.3) with the fabales_odb12 database.

### Gene model prediction

Gene model predictions for two accessions of *V. luteola* were accomplished by integrating the results of Braker2 and GeMoMa. For Braker2 prediction, repetitive elements in assembled genome sequences were first soft-masked using RepeatMasker (4.1.2) (https://github.com/rmhubley/RepeatMasker) with the accession-specific repeat sequence library generated by RepeatModeler (2.0.3) (https://github.com/Dfam-consortium/RepeatModeler). Next, an ab initio gene model prediction was performed on the soft-masked genome with Braker2 (2.1.5) (Brůna *et al*. 2021) and GeneMark (Brůna *et al*. 2020) using the “odb10_plant” protein database from OrthoDB (Kriventseva *et al*. 2019) for protein homology information. Gene models showing homology with the database were selected using a script (https://github.com/Gaius-Augustus/BRAKER/tree/report/scripts/predictionAnalysis) and utilized for subsequent analyses. For GeMoMa analysis, de novo transcriptome assembly was first conducted with Trinity (2.13.2) (Grabherr et al. 2011) using quality-filtered RNA-seq reads from root and leaf tissues of the *V. luteola* tolerant accession. Default parameters were applied for assembly, and the resulting assembly was evaluated using BUSCO (5.2.2) (Manni *et al*. 2021) against “fabale_odb10”, confirming assembly quality (94.3% of conserved orthologs were identified and classified as complete). Subsequently, the same RNA-seq reads were mapped to the de novo transcriptome assembly using HISAT2 (2.2.1) (Kim *et al*. 2019), and mapped reads were retrieved using samtools (1.15.1) (Danecek *et al*. 2021). These reads were utilized for intron and UTR prediction during the GeMoMa process. Homology-based gene prediction was performed by transferring the high-quality gene model of *V. marina* (Naito *et al*. 2022) using GeMoMa (1.8) (Keilwagen *et al*. 2019). The Braker2 prediction was integrated with the GeMoMa results using gffcompare (0.12.6) (Pertea and Pertea 2020). Gene models with the class code “u” were added to the GeMoMa result. Functional gene annotation was carried out on the resulting gene models using EnTAP (0.10.8) (Hart *et al*. 2020). TrEMBL and SwissProt (https://www.uniprot.org/) and NCBI plant RefSeq (https://ncbi.nlm.nih.gov/refseq/) were employed as databases for homology search.

Unannotated gene model entries were removed from the final gene models (31,191 genes for the tolerant accession, 33,814 genes for the sensitive accession). Finally, gene models were renamed using GeMoMa annotation finalizer. The quality of the final gene models was assessed using BUSCO (5.8.3) against fabales_odb12, and 97.8% and 98.5% single-copy orthologs were identified and classified as complete for tolerant and sensitive accessions, respectively.

### Determining the QTL region in the genome of the tolerant accession

The genome of the tolerant accession (Noda *et al*. 2025) was used for this analysis. Chankaew et al. (2014) identified the major QTL related to salt tolerance, located at 101.6 cM of chromosome 1 between the marker CEDG087 and cp06039. We performed BLASTN (Camacho *et al*. 2009) using the marker sequence as the query and a custom database, created from the FASTA file of chromosome 1, as the subject.

### Differential expression analysis

Root transcriptome data (Noda *et al*. 2025) of the tolerant and sensitive accessions was used for transcriptome analysis. Read counts for each gene were estimated using Salmon (1.10.3) (Patro *et al*. 2017) with default parameters. In this process, the transcriptome assembly of the tolerant accession was used as the reference. The obtained count data was used for differential expression analysis using edgeR (4.0.11) (Robinson *et al*. 2010). We followed the protocol of the case study (Fu *et al*. 2015) in the edgeR user’s guide. We compared the expression levels of each gene in the tolerant accession under salt stress and the sensitive accession under salt stress at the same time point. ANOVA-like tests were not performed; instead, differential expression tests were conducted separately for each group.

Genes meeting the following criteria in at least two of the four time points were selected as differentially expressed genes (DEGs): logFC > 0 (upregulated in the tolerant accession) with P-value < 0.05 and FDR < 0.05, or logFC < 0 (downregulated in the tolerant accession) with P-value < 0.05 and FDR < 0.05.

### Gene clustering with self-organizing maps for candidate gene selection

DEGs were clustered with self-organizing maps (SOM) (Wehrens and Buydens 2007), following the protocol described in this script (https://iamciera.github.io/SOMexample/).

Several map sizes, including 4×4, 4×5, and 6×6, were tested to determine the optimal map size. The optimal map size was selected by considering the clarity of the generated clusters.

Clusters showing high expression in the tolerant accession under salt stress and low expression in the sensitive accession under all conditions were used to identify candidate genes.

### Homology search and amino acid comparison

A Homology search was performed with DIAMOND (Buchfink *et al*. 2021) blastp with option “--max-target-seqs 1” and “--evalue 1e-5”. Protein sequences of *A. thaliana* were obtained from The Arabidopsis Information Resource (TAIR) (https://www.arabidopsis.org/download_files/Proteins/Araport11_protein_lists/Araport11_pep_20220914_representative_gene_model.gz). Gene Ontology (GO) annotations of *Arabidopsis thaliana* were downloaded from TAIR (https://www.arabidopsis.org/download_files/GO_and_PO_Annotations/Gene_Ontology_Annotations/ATH_GO_GOSLIM.txt.gz), and genes related to salt stress or abscisic

acid (ABA) were identified based on GO terms or gene descriptions. Specifically, GO terms containing “salinity,” “salt,” or “sodium” were classified as related to salt stress (Supplemental Table 1). Those containing “abscisic acid” were classified as related to abscisic acid (Supplemental Table 2).

Amino acid sequences of selected genes were aligned using ClustalW (2.1) (Larkin *et al*. 2007) with the slow/accurate pairwise alignment option. Additionally, some sequences were analyzed with InterPro (Paysan-Lafosse *et al*. 2023) to detect domain positions using the default options on the website.

### Promoter analysis of four candidate genes

We extracted 2.5 kb regions upstream of the transcription start site (TSS) of candidate genes from both tolerant and sensitive accessions. The extracted promoter sequences were analyzed using FIMO in the MEME suite (Grant *et al*. 2011) with Arabidopsis DAP motifs (O’Malley *et al*. 2016) to identify potential binding motifs. The identified motifs were then compared between the accessions to identify those unique to either the tolerant or sensitive accession.

Subsequently, the promoters of cluster 14 genes and other genes (used as a control) from the tolerant accession were analyzed using FIMO, as described above. The number of genes containing tolerant-specific motifs—identified in candidate gene promoters—was counted for each group. A binomial distribution test was conducted to compare the proportions of motif-containing genes between cluster 14 and the control group. Additionally, the relative frequencies of motifs significantly enriched in cluster 14 were analyzed across the promoter regions and compared with those in the control group.

## Results

### Salt tolerance test of grafted plants

To elucidate the importance of the root and shoot on salt tolerance, we generated grafted plants using the tolerant (T) and sensitive (S) accessions. The salt tolerance test of grafted plants showed that T/T (scion/rootstock) plants had the lowest wilt score, followed by S/T plants and T/S plants, while S/S plants had the highest wilt score (Fig. 1). S/S plants almost completely wilted only five days after salt treatment. Initially, T/S plants maintained significantly lower wilt scores than S/S plants, however, they got high wilt scores 15 days after salt treatment. In contrast, T/T and S/T plants maintained low wilt scores even 15 days after salt treatment. These results suggest that the root of the tolerant accession plays a critical role in conferring salt tolerance in *V. luteola*.

**Figure 1.**
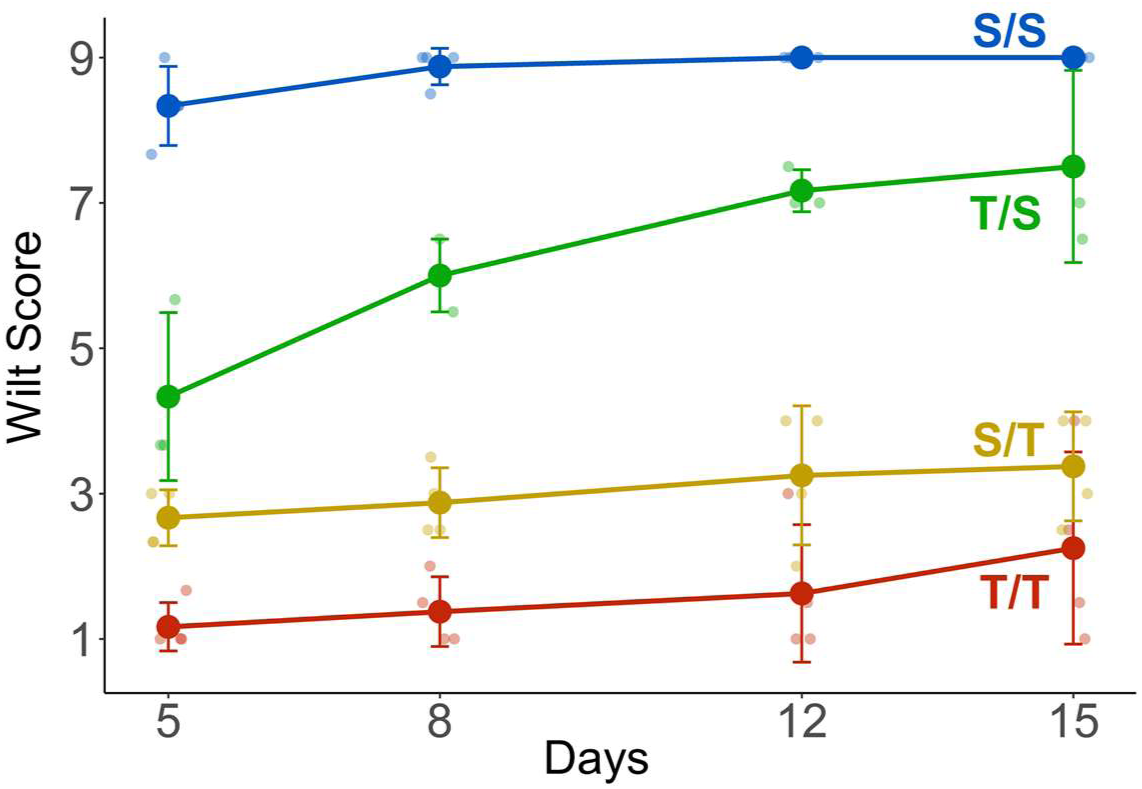
The salt tolerance test of grafted plants under 200 mM NaCl. Small, pale-colored points represent the wilt scores of individual plants. The larger points indicate mean values for each line. Error bars indicate standard deviations. T/T, S/T, S/S, and T/S indicate the scion/rootstock combinations. T and S represent the tolerant and sensitive accessions, respectively.

### Na^+^ and K^+^ allocation in the two accessions

We examined Na^+^ and K^+^ concentrations in whole shoots and roots under both salt and control conditions to determine the role of the root in Na^+^/K^+^ homeostasis.

Under control conditions, Na^+^ concentrations were quite low compared to salt conditions, but the root of the tolerant accession accumulated the most Na^+^ (Fig. 2A). K^+^ concentrations in the shoot were not significantly different between the two accessions, with both maintaining approximately 45 mg/gDW (Fig. 2B). Similar to Na^+^ concentration, Na^+^/K^+^ ratios were quite low under control conditions, but the root of the tolerant accession had the highest ratio (Fig. 2C).

**Figure 2.**
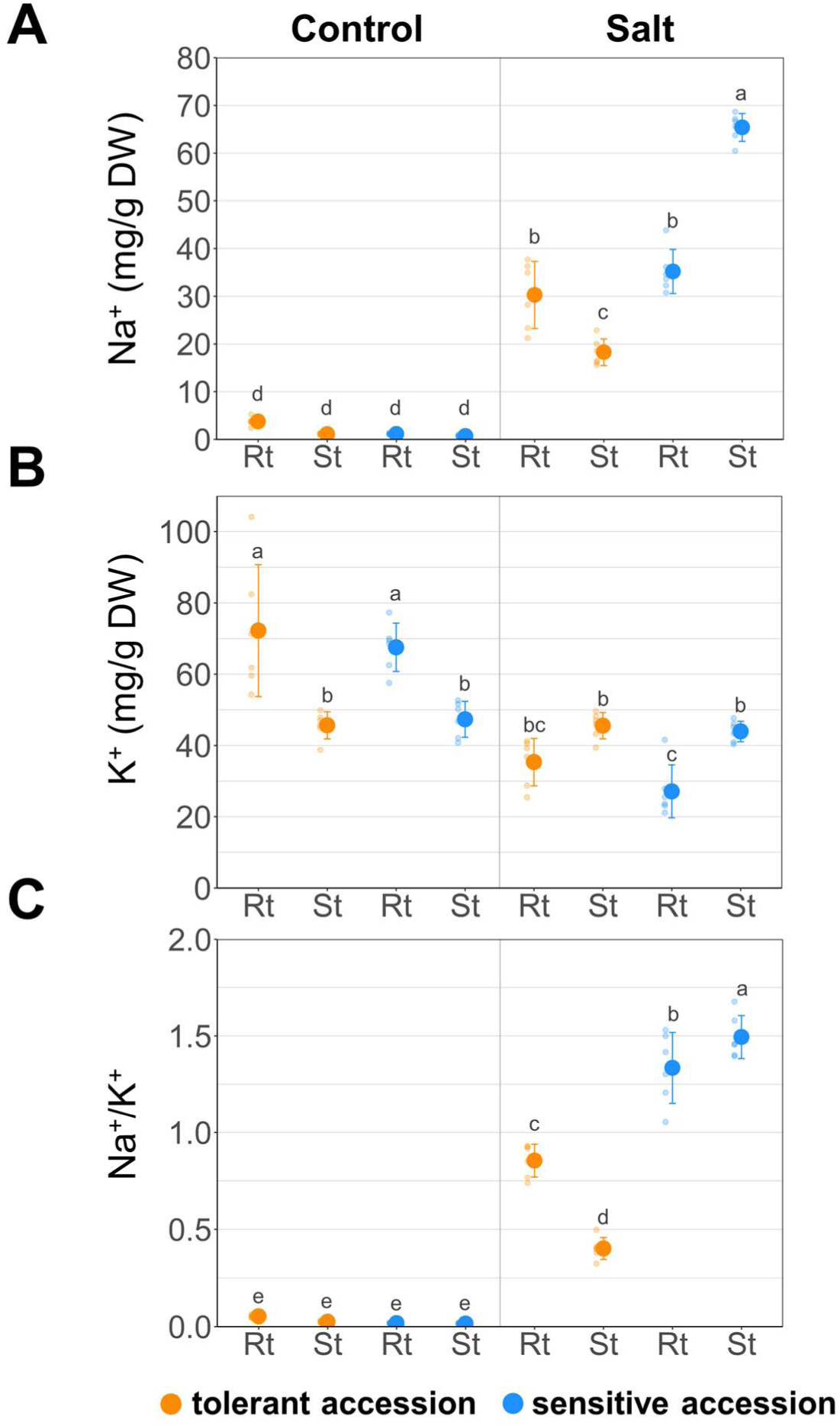
Na^+^ and K^+^ concentrations and the Na⁺/K⁺ ratio in roots (Rt) and shoots (St) of tolerant and sensitive accessions under control and salt (200 mM NaCl) treatment for 3 days. (A) Na^+^ concentration (mg/g DW). (B) K^+^ concentration (mg/g DW). (C) Na^+^/K^+^ ratio. The small, pale-colored points represent values from individual root and shoot samples (n = 6). Larger points and error bars represent the mean and standard deviation, respectively. Different letters above the plots indicate significant differences determined by the Tukey’s HSD test (p < 0.05).

Under salt stress, Na^+^ concentrations in the shoot were significantly higher in the sensitive accession compared to the tolerant accession. In contrast, Na^+^ concentration in the root under salt stress did not differ significantly between the two accessions. Notably, the tolerant accession showed higher Na^+^ concentration in the root than the shoot, whereas the sensitive accession exhibited the opposite pattern, with higher Na^+^ concentrations in the shoot than in the root (Fig. 2D). Shoot K^+^ concentrations did not differ significantly between the two accessions, with both maintaining approximately 45 mg/g DW, similar to the control conditions. The root K^+^ concentration was slightly lower in the sensitive accession than in the tolerant one but not significant (Fig, 2E). The Na^+^/K^+^ ratio was considerably lower in the tolerant accession compared to the sensitive accession, particularly in the shoot (Fig. 2F). These results suggest that the tolerant accession maintains a lower Na⁺/K⁺ ratio in the shoot by suppressing Na⁺ accumulation in the shoot.

### Na^+^ and K^+^ allocation in the grafted plants

To test whether Na^+^ accumulation in the shoot is controlled by the root or the shoot, we analyzed Na^+^ and K^+^ allocation in grafted plants under salt stress.

Grafted plants with the same rootstock exhibited similar trends: T/T and S/T showed similar trends, while S/S and T/S also showed similar trends (Fig. 3). For example, shoot Na^+^ concentrations were significantly lower in T/T and S/T than in S/S and T/S, although root Na^+^ concentrations showed no substantial differences among T/T, S/T, S/S, and T/S. Both T/T and S/T showed lower Na^+^ concentrations in the shoot than in the root. In contrast, S/S and T/S exhibited higher Na^+^ concentrations in the shoot than in the root.

**Figure 3.**
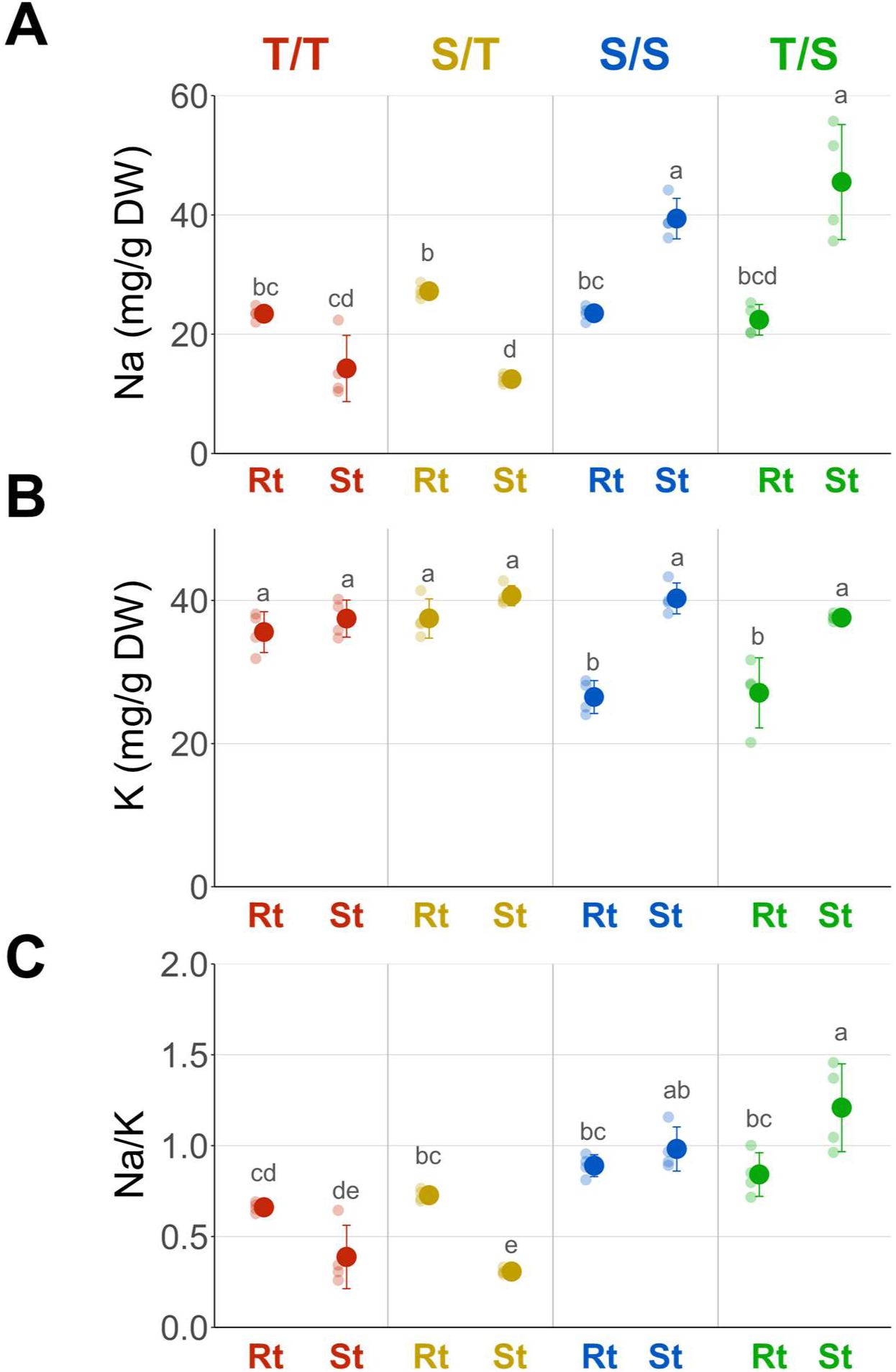
Na⁺ and K⁺ concentrations and the Na⁺/K⁺ ratio in roots (Rt) and shoots (St) of grafted plants under salt (200 mM NaCl) treatment for three days. (A) Na^+^ concentration (mg/g DW). (B) K^+^ concentration (mg/g DW). (C) Na^+^/K^+^ ratio. Small, pale-colored points represent values from individual root and shoot samples (n = 4). Larger points and error bars indicate the mean and standard deviation, respectively. Different letters above the plots indicate significant differences determined by the Tukey’s HSD test (p < 0.05). T/T, S/T, S/S, and T/S indicate the scion/rootstock combinations. T and S represent the tolerant and the sensitive accessions, respectively.

Shoot K^+^ concentration did not differ among T/T, S/T, S/S, and T/S, although the root K^+^ concentration was lower in S/S and T/S than in T/T and S/T.

Shoot Na^+^/K^+^ ratio was significantly lower in T/T and S/T compared to S/S and T/S. Moreover, T/T and S/T showed a lower Na^+^/K^+^ ratio in the shoot than in the root. In contrast, S/S and T/S showed a higher Na^+^/K^+^ ratio in the shoot than in the root. These results suggest that the root plays a primary role in suppressing Na^+^ transport to the shoot.

### Genome assembly and determining the physical position of the QTL region

For QTL localization and comparison of amino acid and promoter sequences between the two accessions, we re-annotated the genome of the tolerant accession (Noda *et al*. 2025) and newly assembled the genome of the sensitive accession.

We sequenced the whole genome of the sensitive accession using Oxford Nanopore. The draft genome assembly contained 496.2 Mbp in 210 contigs with an N50 length of 11.5 Mbp. These contigs were further scaffolded into 11 scaffolds (Supplemental Tables 3 and 4). BUSCO analysis (v5.8.3, fabales_odb12) of the assembly showed 98.4% completeness in the sensitive accession. For comparison, we re-analyzed the genome assembly of the tolerant accession (Noda et al., 2025) using the same version and database, which showed 98.5% completeness. Gene annotation for the two accessions predicted 31,191 genes in the tolerant accession and 33,814 genes in the sensitive accession, covering 97.8% and 98.5% of BUSCO genes (v5.8.3, fabales_odb12), respectively.

Subsequently, we identified the genomic region corresponding to the major QTL for salt tolerance, previously identified by Chankaew et al. (2014), in the tolerant accession genome. The major QTL region ranged from 57,314,479 bp to 62,876,770 bp on chromosome 1 and contained 557 genes.

### Gene expression analysis and the candidate genes

We performed gene expression analysis under salt stress to further narrow down candidate genes.

First, we identified differentially expressed genes (DEGs) between the two accessions under salt stress using the root transcriptome data (Noda *et al*. 2025). The number of genes upregulated in the tolerant accession was 1,709 in the first daytime (1D), 4,021 in the first nighttime (1N), 2,901 in the second daytime (2D), and 3,808 in the second nighttime (2N). The number of genes downregulated in the tolerant accession was 1,652 in 1D, 4,111 in 1N, 2,484 in 2D, and 3,915 in 2N (Supplemental Fig. 2). A total of 6,917 genes detected at least 2 time points were used for further clustering.

SOM-clustering divided the DEGs into 16 clusters (Supplemental Fig. 3). Among the 16 clusters, we focused on three clusters— 9, 13, and 14—that showed high expression under salt stress in the tolerant accession (Fig. 4A). These clusters contained 375 genes (cluster 9), 526 genes (cluster 13), and 401 genes (cluster 14). In the three candidate clusters, 27 genes were in the QTL region (Supplemental Fig. 4, Supplemental Table 5). Among those, 25 genes had Arabidopsis homologs, whereas the remaining two genes contained no detectable domain (Supplemental Table 5). Four genes were selected for further analysis by GO terms (Supplemental Tables 1 and 2) or gene function related to salt tolerance (Fig. 4B).

**Figure 4.**
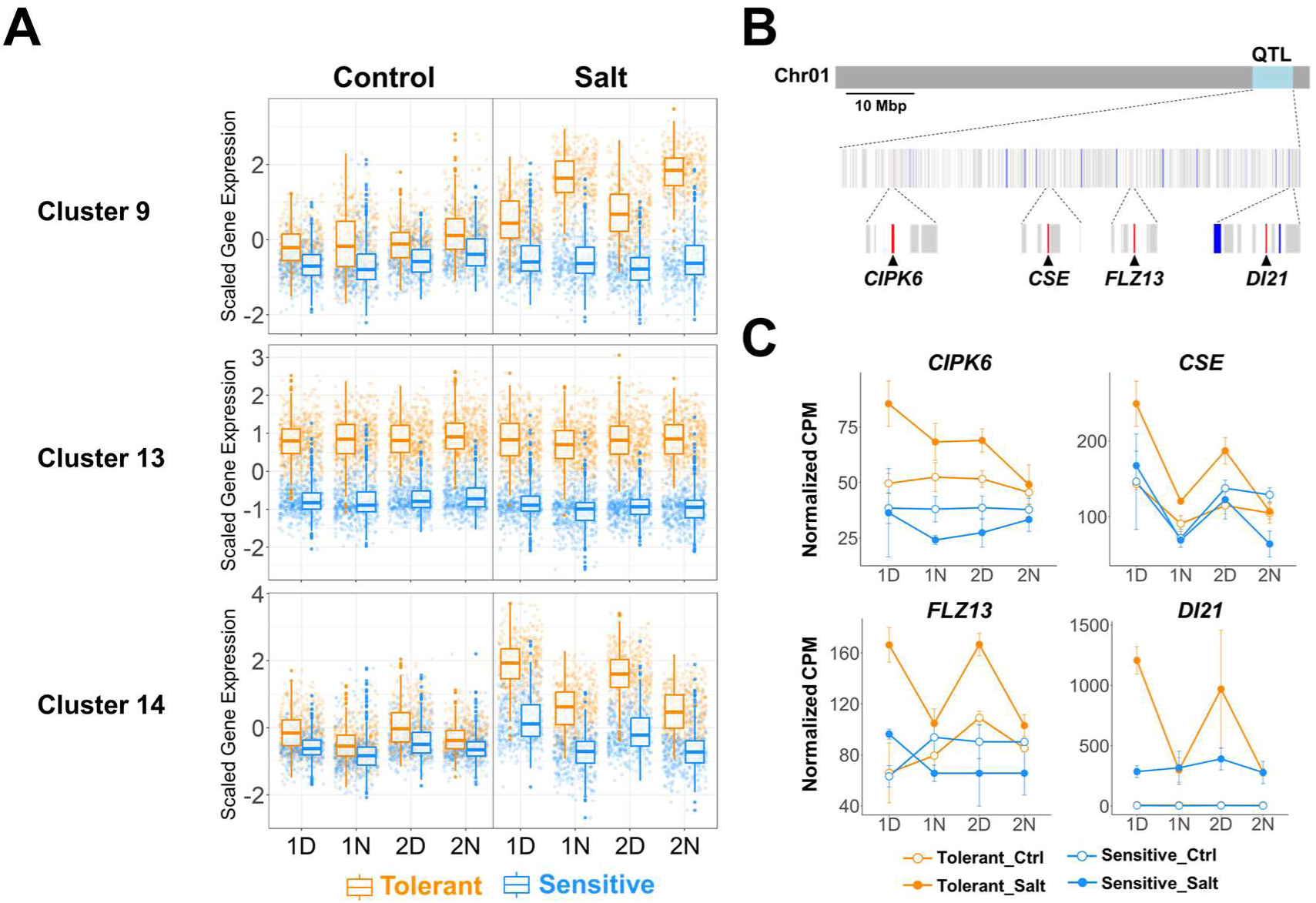
Expression patterns of candidate clusters and the candidate genes. (A) Expression patterns of the three candidate clusters that are up-regulated in the tolerant accession under salt stress. Tolerant and Sensitive indicate the tolerant and sensitive accessions, respectively. Control and Salt indicate control (0 mM NaCl) and salt stress (200 mM NaCl) conditions, respectively. 1D, 1N, 2D, and 2N represent 1st daytime, 1st nighttime, 2nd daytime, and 2nd nighttime respectively. (B) Genomic positions of genes within the QTL region and candidate clusters. The light blue region indicates the major QTL (*Saltol1.1*) region associated with salt tolerance. Vertical bars represent individual genes within the QTL region. Genes belonging to the candidate clusters are shown in blue, while the four candidate genes (CIPK6, CSE, FLZ13, and DI21) are shown in red. Other genes in the region are shown in gray. *CIPK6*, *FLZ13*, *CSE*, and *DI21* indicate *Viglu256296.01G0396300.1*, *Viglu256296.01G0413000.1*, *Viglu256296.01G0424800.1*, and *Viglu256296.01G0445600.1*, respectively. (C) Expression levels of the four candidate genes. The dots and error bars indicate the mean and standard deviation of the biological replicates of normalized CPM, respectively. Tolerant and Sensitive indicate the tolerant accession and the sensitive accession, respectively. Ctrl and Salt indicate control and salt stress conditions, respectively. Tolerant_Ctrl is not visible because it overlaps with Sensitive_Ctrl due to identical values in *DI21*.

The four genes were *Viglu256296.01G0396300.1*, homologous to *CBL-interacting protein kinase 6* (*CIPK6*), with the GO term “response to salt stress”; *Viglu256296.01G0413000.1*, homologous to AT1G52760.1 *Caffeoyl Shikimate Esterase* (*CSE*), reported to be involved in lignin biosynthesis; *Viglu256296.01G0424800.1*, homologous to *AT1G74940.1 FCS-LIKE ZINC FINGER PROTEIN 13* (*FLZ13*), with the GO term “response to salt stress”; and *Viglu256296.01G0445600.1*, homologous to *AT4G15910.1 drought-induced 21* (*DI21*), with the GO term “response to abscisic acid”. All four genes belonged to cluster 14 (Fig. 4A) and were highly expressed in the tolerant accession under salt stress, particularly during the daytime (Fig. 4C).

A comparison of the amino acid sequences of the tolerant accession and the sensitive accession showed that CIPK6 and FLZ13 had insertion-deletion mutations and substitutions (Supplemental Figs. 5 and 6), whereas DI21 and CSE were entirely identical (Supplemental Fig. 7). However, the mutations were not within the conserved domains (Supplemental Figs. 5 and 6). These findings suggest that the protein functions of the four candidate genes are highly conserved between the two accessions, and that their differential expression may contribute to salt tolerance in the tolerant accession.

### Promoter analysis on CIPK6, CSE, FLZ13, and DI21

We next examined the promoter regions to explore potential *cis*-regulatory differences. *CIPK6*, *CSE, FLZ13*, and *DI21* are located within the major QTL region, but it remains unclear whether their expression is regulated by *cis*-regulatory elements in the QTL. To investigate this, we analyzed promoter sequences for differences in transcription factor binding sites (TFBSs) between the two accessions. Because the relative distance from the TSS is known to influence gene expression in plants (Voichek *et al*. 2024), we also examined the positional distribution of TFBSs.

First, promoter regions of the four genes were analyzed through motif scanning to identify potential binding motifs. As a result, 12, 20, 11, and 3 distinct motifs unique to tolerant accession were identified in the promoters of *CIPK6*, *CSE*, *FLZ13*, and *DI21,* respectively (Fig. 5A, Supplemental Fig. 8).

**Figure 5.**
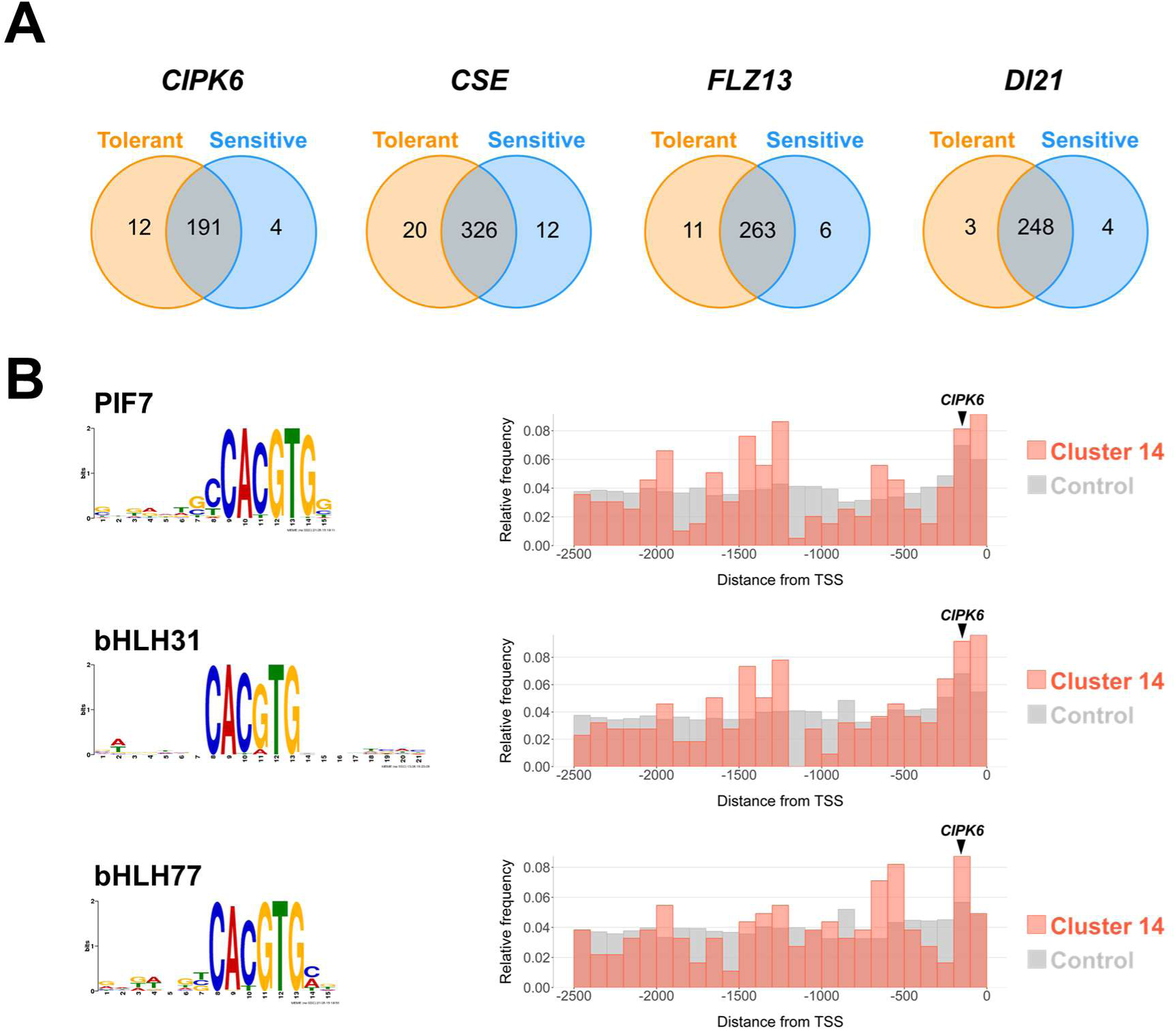
Promoter analysis of the candidate genes. (A) Number of unique and shared motifs of the promoters (2.5 kb upstream from TSS) in the two accessions. (B) Transcription factor binding motifs unique to the tolerant accession in the *CIPK6* promoter, enriched in cluster 14 promoters compared to control, and are located at positions corresponding to motif-enriched regions in cluster 14 promoters. Pictograms of the motifs and relative frequencies of these motifs across the promoters of cluster 14 and control groups are shown. Black arrows indicate the positions of the motifs on the *CIPK6* promoter.

To investigate whether these motifs are associated with gene expression patterns, we also analyzed the promoters of cluster 14 genes and other genes (used as control). The 3, 11, 3, and 1 motifs in *CIPK6*, *CSE*, *FLZ13*, and *DI21*, respectively, were significantly enriched in cluster 14 compared to control genes (Supplemental Table 6). In addition, investigation of the position from TSS revealed that binding motifs for PIF7, bHLH31, and bHLH77 were enriched within 200 bp upstream of the TSS in the promoters of cluster 14, corresponding to the location of the *CIPK6* promoter (Fig. 5B). However, no such motifs were detected in the promoters of *CSE*, *FLZ13*, and *DI21* at positions corresponding to the motif-enriched regions in cluster 14 promoters (Supplemental Fig. 9).

Furthermore, we investigated whether the transcription factors PIF7, bHLH31, and bHLH77 also belonged to cluster 14. As a result, we found that *Viglu256296.01G0358900.3*, a homolog of bHLH31 (BPEp, *AT1G59640.2*), was present in cluster 14 and showed similar expression patterns with *CIPK6* (Supplemental Fig. 10A). Amino acid comparison showed that the bHLH_AtBPE_like domain region is highly conserved between Viglu256296.01G0358900.3 and AT1G59640.2 (Supplemental Fig. 10B).

## Discussion

In this study, we demonstrated that suppressing Na^+^ transport to the shoot plays a major role in the salt tolerance of the tolerant accession of *V. luteola*. By integrating root transcriptome data and QTL, we narrowed down the candidate genes for the luteola’s salt tolerance, including functionally important genes such as *CIPK6*, *CSE*, *FLZ13*, and *DI21*.

The promoter analysis revealed that a *cis*-element within 200 bp upstream of *CIPK6* is associated with induced expression under salt stress.

### Suppressing Na^+^ transport to the shoot is a primary factor in salt tolerance

Given that S/T plants showed lower wilt scores than T/S plants, the root should play a more important role in the salt tolerance of *V. luteola* (Fig. 1). This, together with the fact that grafted plants with rootstock of the tolerant accession significantly suppressed Na^+^ allocation to the shoot (Fig. 3A), indicates that the root of tolerant accession has higher ability to block Na^+^ transport. In contrast, the grafted plants did not show significant differences in K^+^ allocation to the shoot (Fig. 3B). Thus, the suppression of Na^+^ transport in the tolerant accession does not rely on increased K^+^ uptake. This indicates that the mechanism of *V. luteola* is different from many other salt-tolerant plants that, in response to salt stress, increase the K^+^ content to prevent Na^+^ uptake (Kumari *et al*. 2021; Ito *et al*. 2024). Previously we have elucidated that the tolerant accession of *V. luteola* suppresses Na^+^ transport by lignification of Casparian strip and excludes Na^+^ from the root by increasing expression of *Salt Overly Sensitive 1* (*SOS1*) that encodes a Na^+^/H^+^ antiporter (Noda *et al*. 2025; Wang *et al*. 2025). As the *SOS1* locus is not located in the QTL region, the genes within the QTL could be involved in upregulating the *SOS1* transcription.

In addition, although the T/S and S/S plants allocated a similar amount of Na^+^ to the shoot, the S/S plants showed a higher wilt score (Fig.1, 3). This indicates that the shoot of the tolerant accession can tolerate higher concentrations of tissue Na^+^. Although the contribution is smaller than the mechanism in the root, the difference is still significant and will be worth investigating.

### CIPK6 is the most promising candidate gene for the QTL

The results of this study indicate that *CIPK6* is the most promising candidate gene for salt tolerance. It is located within the previously identified QTL (Chankaew *et al*. 2014), strongly induced by salt stress (Fig. 4C), and has a *cis*-element variation (Fig. 5B) potentially involved in its upregulation by salt stress. *CIPK6* encodes CBL-interacting protein kinase (CIPK), which is activated through its interaction with a calcium sensor, calcineurin B-like protein (CBL) (Batistič and Kudla 2009; He *et al*. 2013). The CBL– CIPK network plays important roles under various stresses (Yu *et al*. 2014). Furthermore, in a previous study of rice, overexpression of *CdtCIPK5*, an ortholog of *AtCIPK6*, enhanced salt tolerance by restricting Na^+^ transport to the shoot (Huang *et al*. 2020). Similarly, the tolerant accession showed low Na^+^ accumulation in the shoot (Fig. 2). Therefore, we consider that *CIPK6* may contribute to suppressing Na^+^ transport in the tolerant accession of *V. luteola*. The conserved amino acid sequences in the functional domains suggest that the CIPK6 protein has similar functions in both accessions (Supplemental Fig. 5), and we hypothesize that the observed difference in expression explains the difference in salt tolerance. The CIPK6 promoter in the tolerant accession harbors binding sites for bHLH transcription factors including PIF7, bHLH31, and bHLH77. These motifs were significantly enriched in the promoters of cluster 14 genes, which were highly expressed under salt stress in the tolerant accession (Fig. 4A). Moreover, these motifs were located within 200 bp upstream from the TSS in *CIPK6*, the same position where they are frequently enriched in cluster 14 promoters (Fig. 5B). Since the distance of transcription factor binding sites (TFBSs) from the TSS is known to influence gene expression (Voichek *et al*. 2024), this positional overlap suggests that these motifs may contribute to the salt-induced expression of CIPK6. Notably, a homolog of bHLH31 showed a similar expression pattern to CIPK6, and the bHLH_AtBPE_like domain was highly conserved between *V. luteola* and *Arabidopsis* (Supplemental Fig. 10). These observations support the possibility that increased expression of bHLH31 in the tolerant accession may drive the higher expression of *CIPK6* under salt stress, though it remains to be tested whether these motifs directly regulate CIPK6 expression. Overall, these results suggest that polymorphisms in the *CIPK6* promoter might be a causal variant for the major QTL.

### Other candidate genes

Besides *CIPK6*, we identified *CSE*, *FLZ13*, and *DI21* as possible candidates. Although no notable differences in *cis*-regulatory motifs associated with gene expression were observed-(Supplemental Fig. 9), these genes are also located within QTL, highly expressed in the tolerant accession, and possess functions potentially related to salt tolerance.

*DI21* would contribute to water uptake under salt stress rather than Na^+^ transport. In previous studies, *DI21* has been up-regulated by progressive drought stress (Gosti *et al*. 1995; Yan *et al*. 2012). Salt stress is known to be associated with osmotic stress (Zhang *et al*. 2006). The tolerant accession of *V. luteola* exhibited higher water uptake under salt stress compared to the sensitive accession, although not statistically significant (Wang *et al*. 2024). While the precise function of *DI21* remains unclear, this gene would contribute to water uptake rather than suppress Na^+^ transport under salt stress.

We consider that *CSE* would contribute to lignification in the Casparian strip formation, thereby preventing Na⁺ transport to the shoot in the root. Previous research has reported that *CSE* is involved in lignin biosynthesis (Vanholme *et al*. 2013). Moreover, disruption of lignification in the root increased Na^+^ transport to the shoot in the tolerant accession of *V. luteola* (Wang *et al*. 2025). Based on these reports, *CSE* is considered to play a role in suppressing Na⁺ transport to the shoot in the root through lignin biosynthesis. *FLZ13* has been reported to interact with *Abscisic acid insensitive5* (*ABI5*) and positively regulate the ABA signaling (Yang *et al*. 2023). ABA signaling is known to play a significant role in drought and salt stress responses (Zhang *et al*. 2006). Although the precise role of *FLZ13* in salt tolerance remains unclear, it is also considered to contribute to the salt stress response.

In addition to the four candidate genes discussed above, the QTL region harbors 23 genes that also exhibited higher expression in the tolerant accession under salt stress (Supplemental Table 5). As we cannot currently exclude the possibility that some of them are also involved in salt tolerance, we need further investigation to identify the responsible gene.

### Conclusions and future perspectives

In conclusion, our findings indicate that suppression of Na^+^ transport to the shoot by the root is essential for salt tolerance in *V. luteola*. Additionally, the high expression of *CIPK6* in the root of the tolerant accession is considered to play a key role in this mechanism. To validate this hypothesis, future complementation studies will be necessary. The candidate genes identified in this study have the potential to contribute to the development of salt-tolerant crops.

## Data Availability Statement

The RNA sequence data is available at the Sequence Read Archive of the NCBI website. The BioProject ID is PRJNA1080052. BioSamples IDs range from SAMN40099594 to SAMN40099689, but only samples containing “R1,” “R2,” and “R3” in the Sample Name were used in this study.

## Acknowledgments

We thank Dr. T. Seiko and Dr. K. Nagasawa (Research Center of Genetic Resources, NARO) for their assistance with wilt scoring during the salt tolerance test. This work was financially supported by SPRING, Japan Science and Technology Agency [grant number JPMJSP2108]; Grant-in-Aid for JSPS Fellows [grant number JP24KJ0550]; and Moonshot R&D Program for Agriculture, Forestry and Fisheries by Cabinet Office [grant number JPJ009237].

## Author Contributions

YI and KN conceived the study.

YI produced grafting plants and performed a salt tolerance test. YI, FW, and KT performed ion measurements.

KI, FW, and KN performed sequencing and genome assembly. TW performed gene model prediction.

YI, FW, and KN analyzed transcriptome data. YI and KN performed promoter analysis.

YI and KN wrote the paper.

