## Supplemental Figures for "Comparative genomics and transcriptomics on salt tolerance of *Vigna luteola*"

### Slide 1
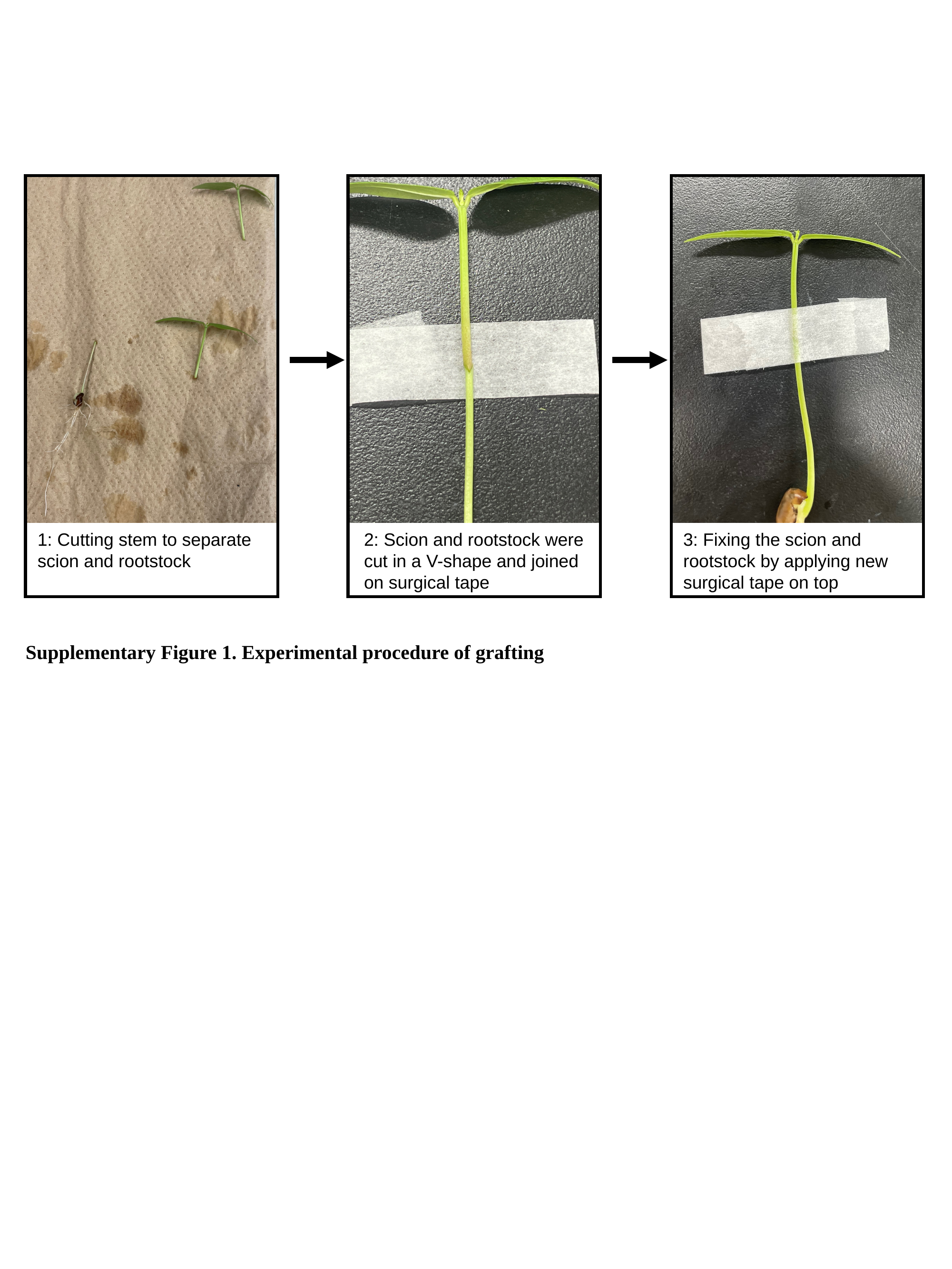

1: Cutting stem to separate scion and rootstock
3: Fixing the scion and rootstock by applying new surgical tape on top
2: Scion and rootstock were cut in a V-shape and joined on surgical tape
Supplementary Figure 1. Experimental procedure of grafting

### Slide 2
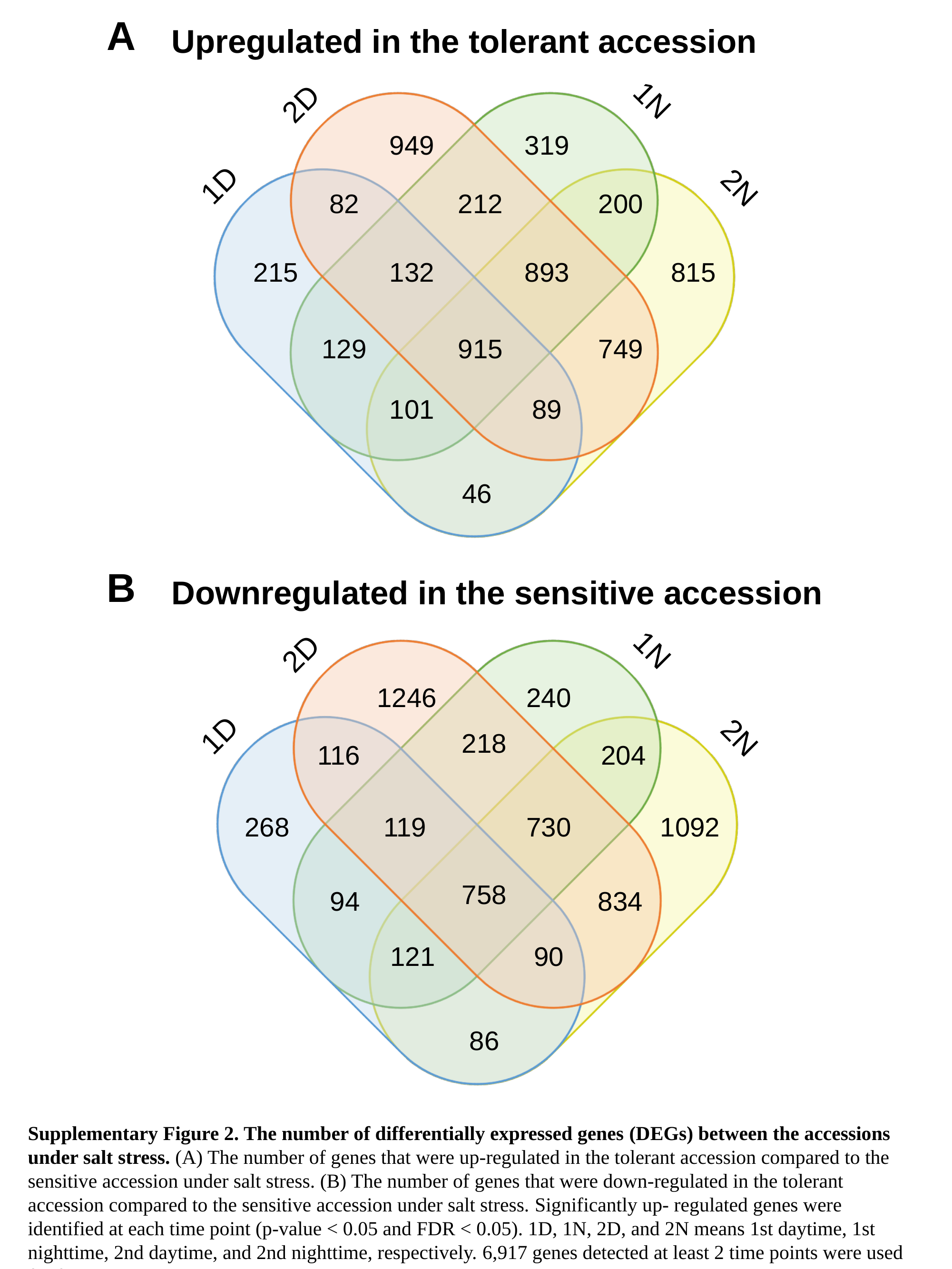

A
Upregulated in the tolerant accession
1N
2N
1D
2D
949
319
82
212
200
215
132
893
815
129
915
749
101
89
46
B
Downregulated in the sensitive accession
1N
2N
1D
2D
1246
240
218
116
204
268
119
730
1092
758
94
834
121
90
86
Supplementary Figure 2. The number of differentially expressed genes (DEGs) between the accessions under salt stress. (A) The number of genes that were up-regulated in the tolerant accession compared to the sensitive accession under salt stress. (B) The number of genes that were down-regulated in the tolerant accession compared to the sensitive accession under salt stress. Significantly up- regulated genes were identified at each time point (p-value < 0.05 and FDR < 0.05). 1D, 1N, 2D, and 2N means 1st daytime, 1st nighttime, 2nd daytime, and 2nd nighttime, respectively. 6,917 genes detected at least 2 time points were used for further clustering.

### Slide 3
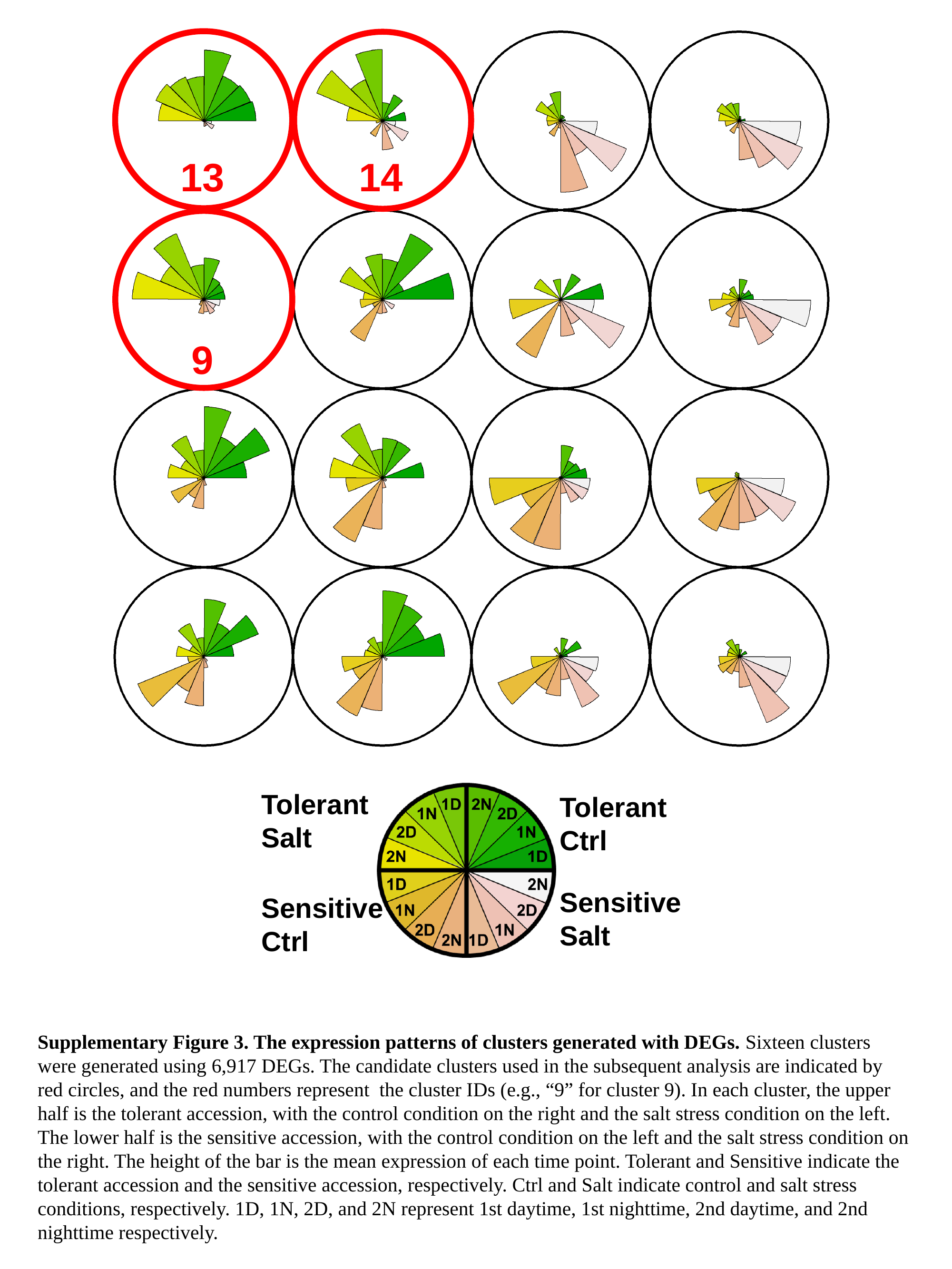

13
14
9
Tolerant
Salt
Tolerant
Ctrl
Sensitive
Salt
Sensitive
Ctrl
Supplementary Figure 3. The expression patterns of clusters generated with DEGs. Sixteen clusters were generated using 6,917 DEGs. The candidate clusters used in the subsequent analysis are indicated by red circles, and the red numbers represent the cluster IDs (e.g., “9” for cluster 9). In each cluster, the upper half is the tolerant accession, with the control condition on the right and the salt stress condition on the left. The lower half is the sensitive accession, with the control condition on the left and the salt stress condition on the right. The height of the bar is the mean expression of each time point. Tolerant and Sensitive indicate the tolerant accession and the sensitive accession, respectively. Ctrl and Salt indicate control and salt stress conditions, respectively. 1D, 1N, 2D, and 2N represent 1st daytime, 1st nighttime, 2nd daytime, and 2nd nighttime respectively.

### Slide 4
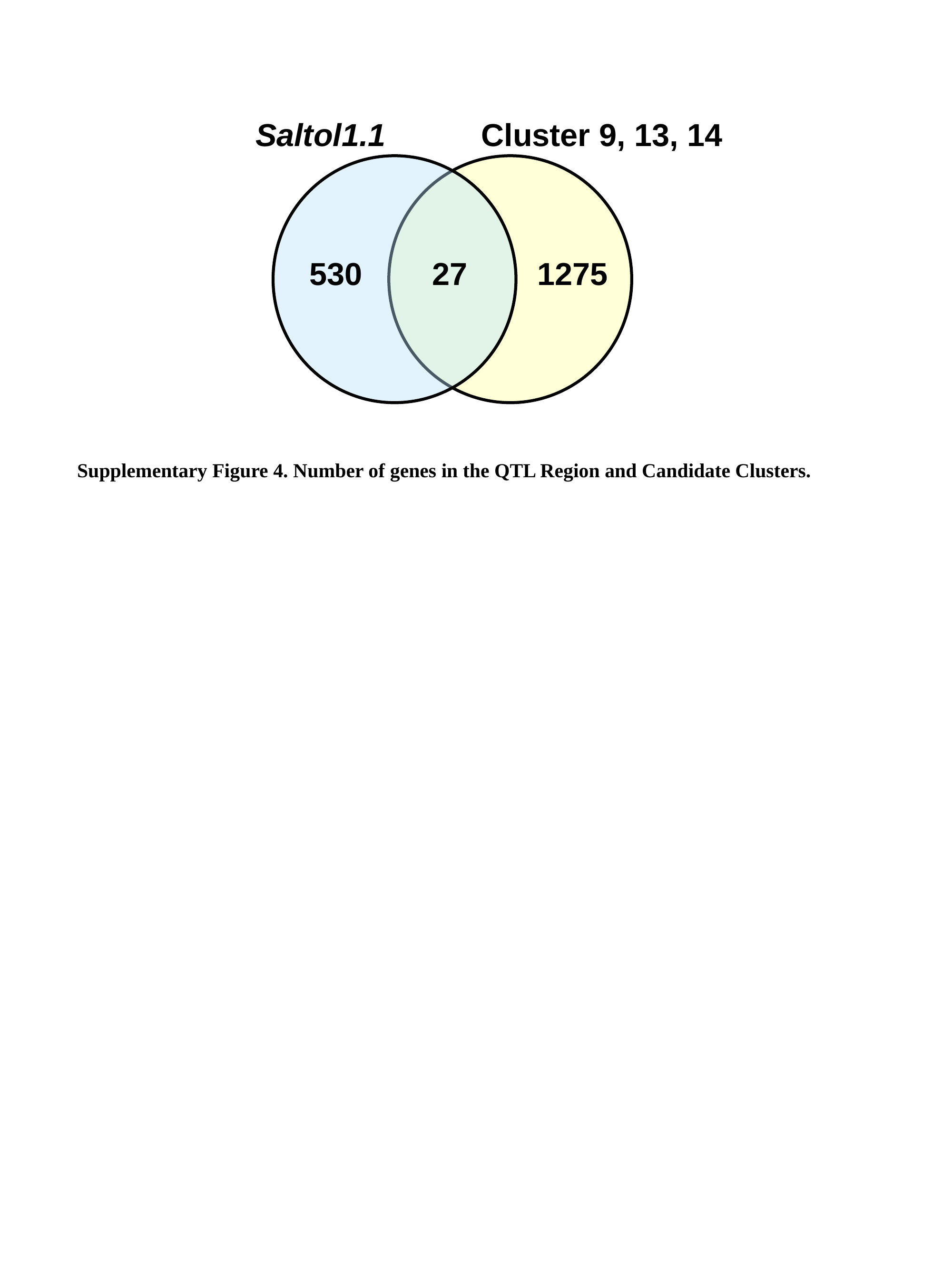

Saltol1.1
Cluster 9, 13, 14
530
27
1275
Supplementary Figure 4. Number of genes in the QTL Region and Candidate Clusters.

### Slide 5
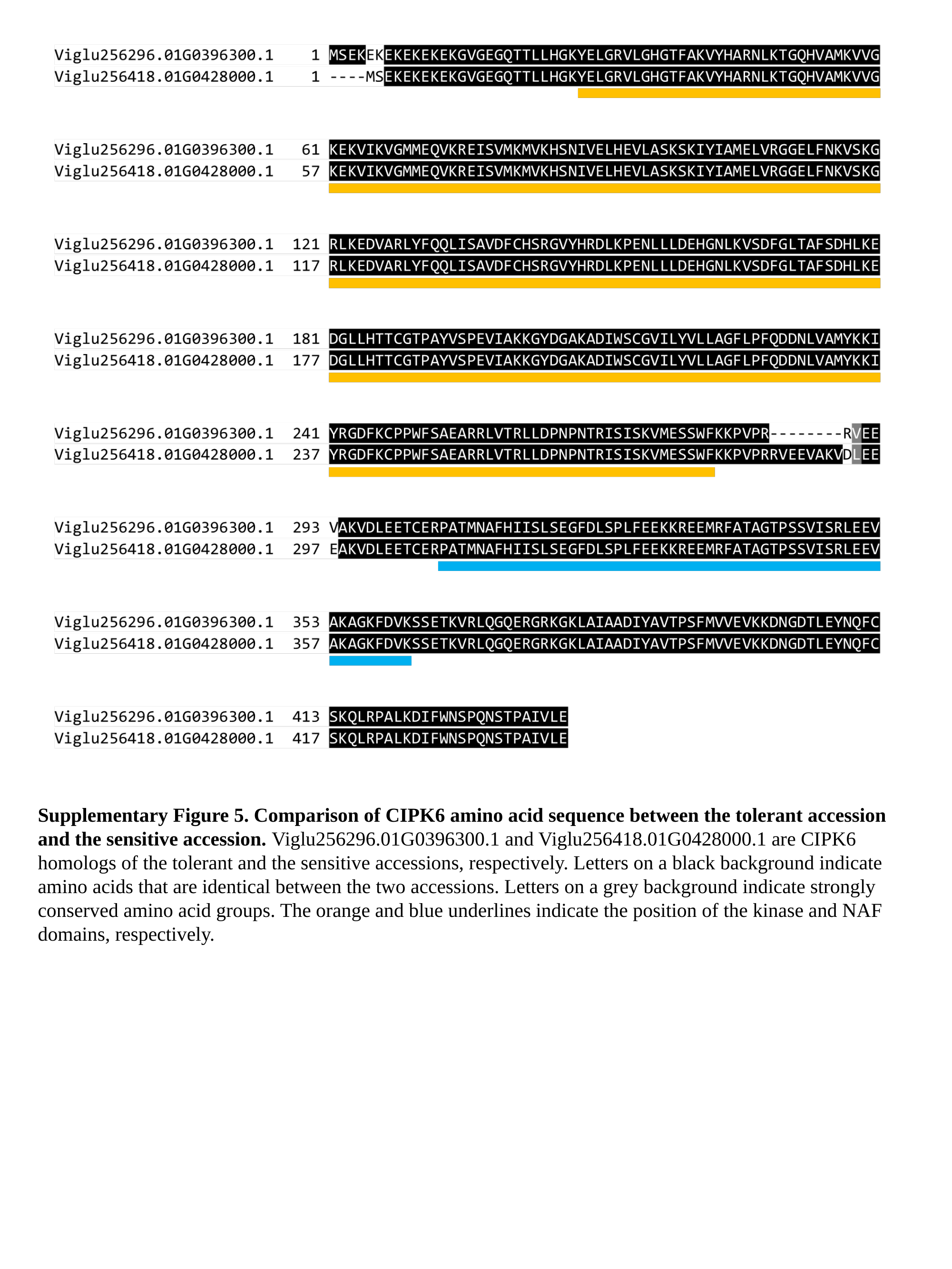

Supplementary Figure 5. Comparison of CIPK6 amino acid sequence between the tolerant accession and the sensitive accession. Viglu256296.01G0396300.1 and Viglu256418.01G0428000.1 are CIPK6 homologs of the tolerant and the sensitive accessions, respectively. Letters on a black background indicate amino acids that are identical between the two accessions. Letters on a grey background indicate strongly conserved amino acid groups. The orange and blue underlines indicate the position of the kinase and NAF domains, respectively.

### Slide 6
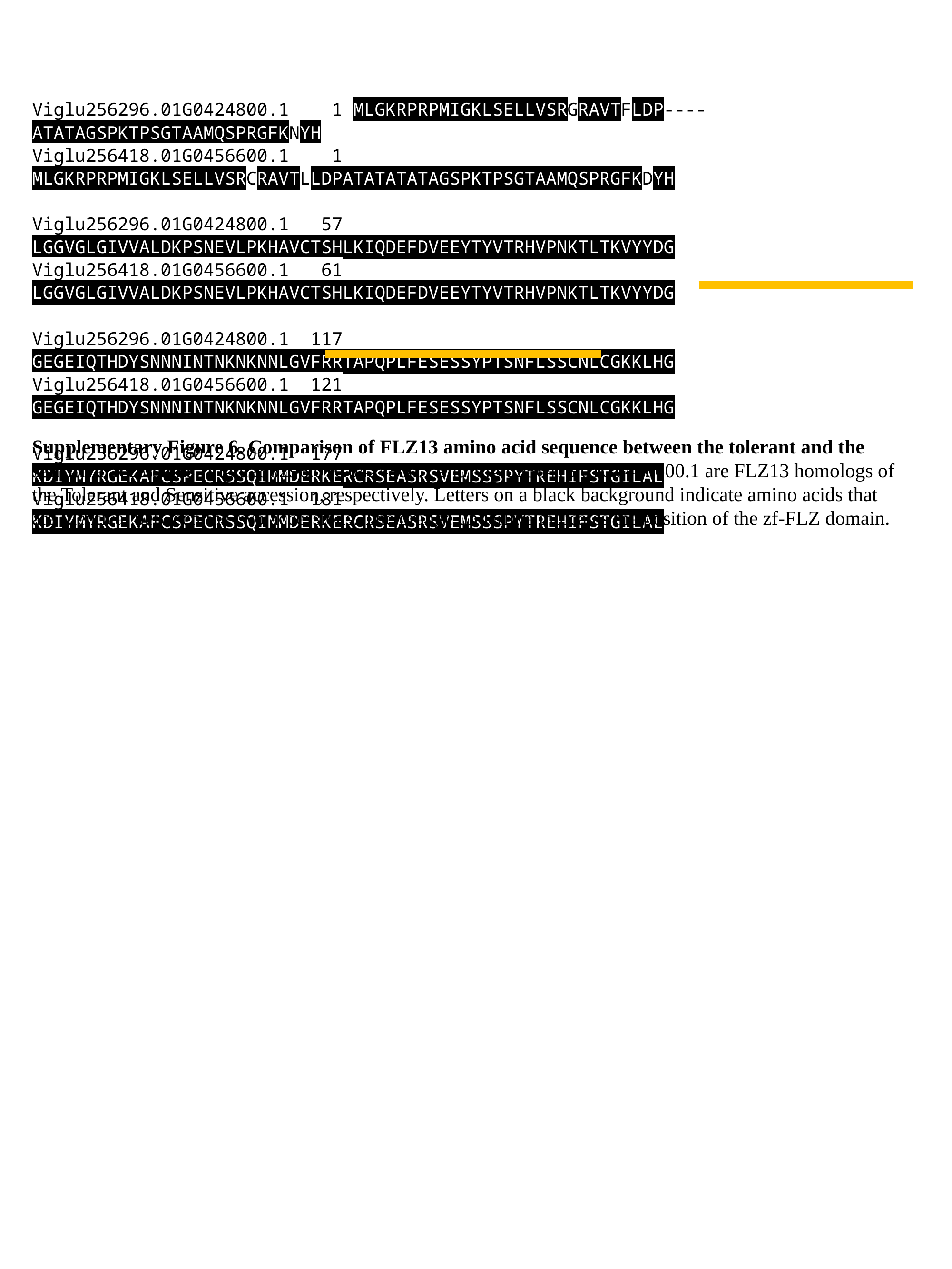

Viglu256296.01G0424800.1 1 MLGKRPRPMIGKLSELLVSRGRAVTFLDP----ATATAGSPKTPSGTAAMQSPRGFKNYH
Viglu256418.01G0456600.1 1 MLGKRPRPMIGKLSELLVSRCRAVTLLDPATATATATAGSPKTPSGTAAMQSPRGFKDYH
Viglu256296.01G0424800.1 57 LGGVGLGIVVALDKPSNEVLPKHAVCTSHLKIQDEFDVEEYTYVTRHVPNKTLTKVYYDG
Viglu256418.01G0456600.1 61 LGGVGLGIVVALDKPSNEVLPKHAVCTSHLKIQDEFDVEEYTYVTRHVPNKTLTKVYYDG
Viglu256296.01G0424800.1 117 GEGEIQTHDYSNNNINTNKNKNNLGVFRRTAPQPLFESESSYPTSNFLSSCNLCGKKLHG
Viglu256418.01G0456600.1 121 GEGEIQTHDYSNNNINTNKNKNNLGVFRRTAPQPLFESESSYPTSNFLSSCNLCGKKLHG
Viglu256296.01G0424800.1 177 KDIYMYRGEKAFCSPECRSSQIMMDERKERCRSEASRSVEMSSSPYTREHIFSTGILAL
Viglu256418.01G0456600.1 181 KDIYMYRGEKAFCSPECRSSQIMMDERKERCRSEASRSVEMSSSPYTREHIFSTGILAL
Supplementary Figure 6. Comparison of FLZ13 amino acid sequence between the tolerant and the sensitive accessions. Viglu256296.01G0424800.1 and Viglu256418.01G0456600.1 are FLZ13 homologs of the Tolerant and Sensitive accession, respectively. Letters on a black background indicate amino acids that are identical between the two accessions. The orange underline indicates the position of the zf-FLZ domain.

### Slide 7
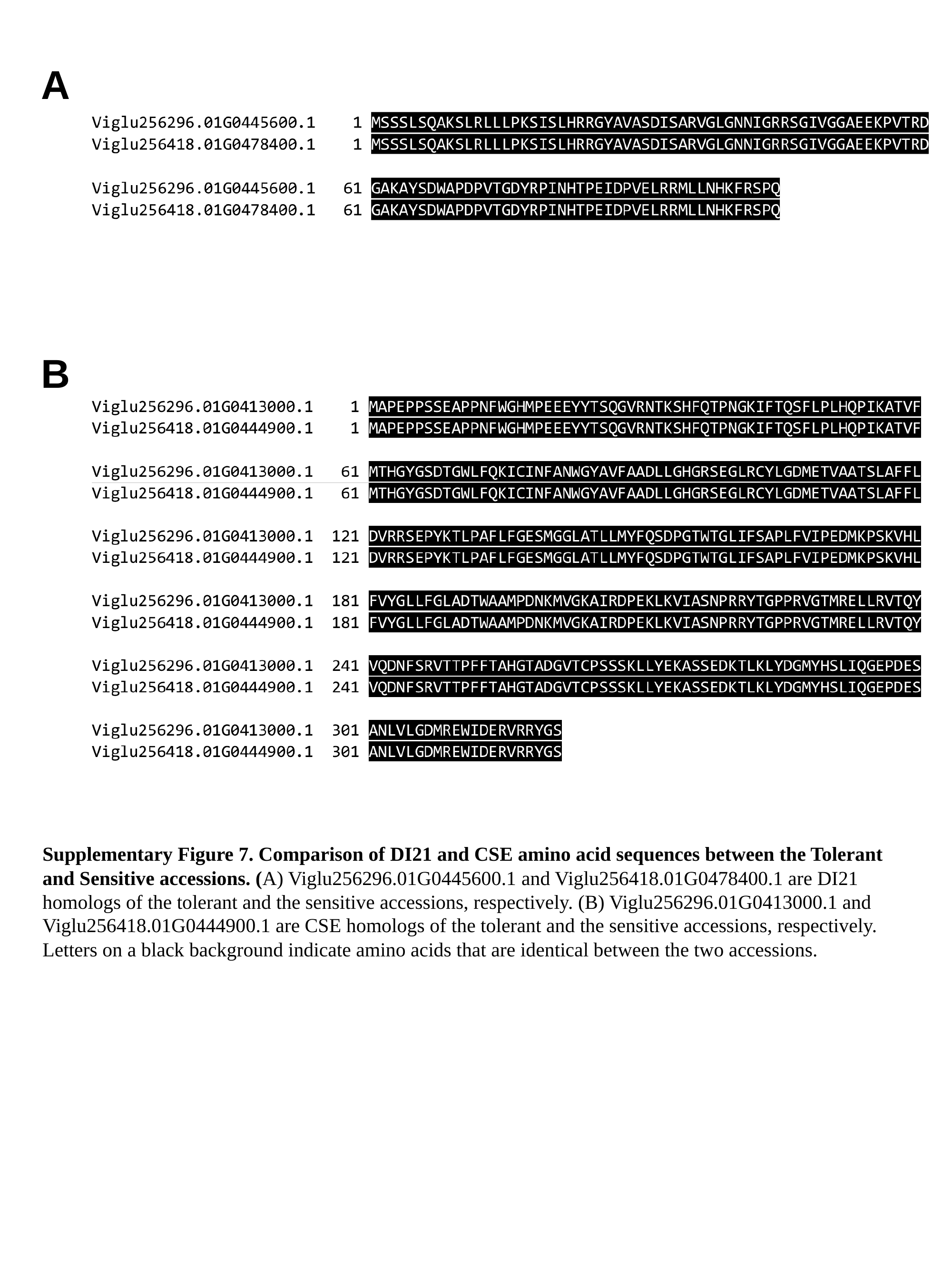

A
B
Supplementary Figure 7. Comparison of DI21 and CSE amino acid sequences between the Tolerant and Sensitive accessions. (A) Viglu256296.01G0445600.1 and Viglu256418.01G0478400.1 are DI21 homologs of the tolerant and the sensitive accessions, respectively. (B) Viglu256296.01G0413000.1 and Viglu256418.01G0444900.1 are CSE homologs of the tolerant and the sensitive accessions, respectively. Letters on a black background indicate amino acids that are identical between the two accessions.

### Slide 8
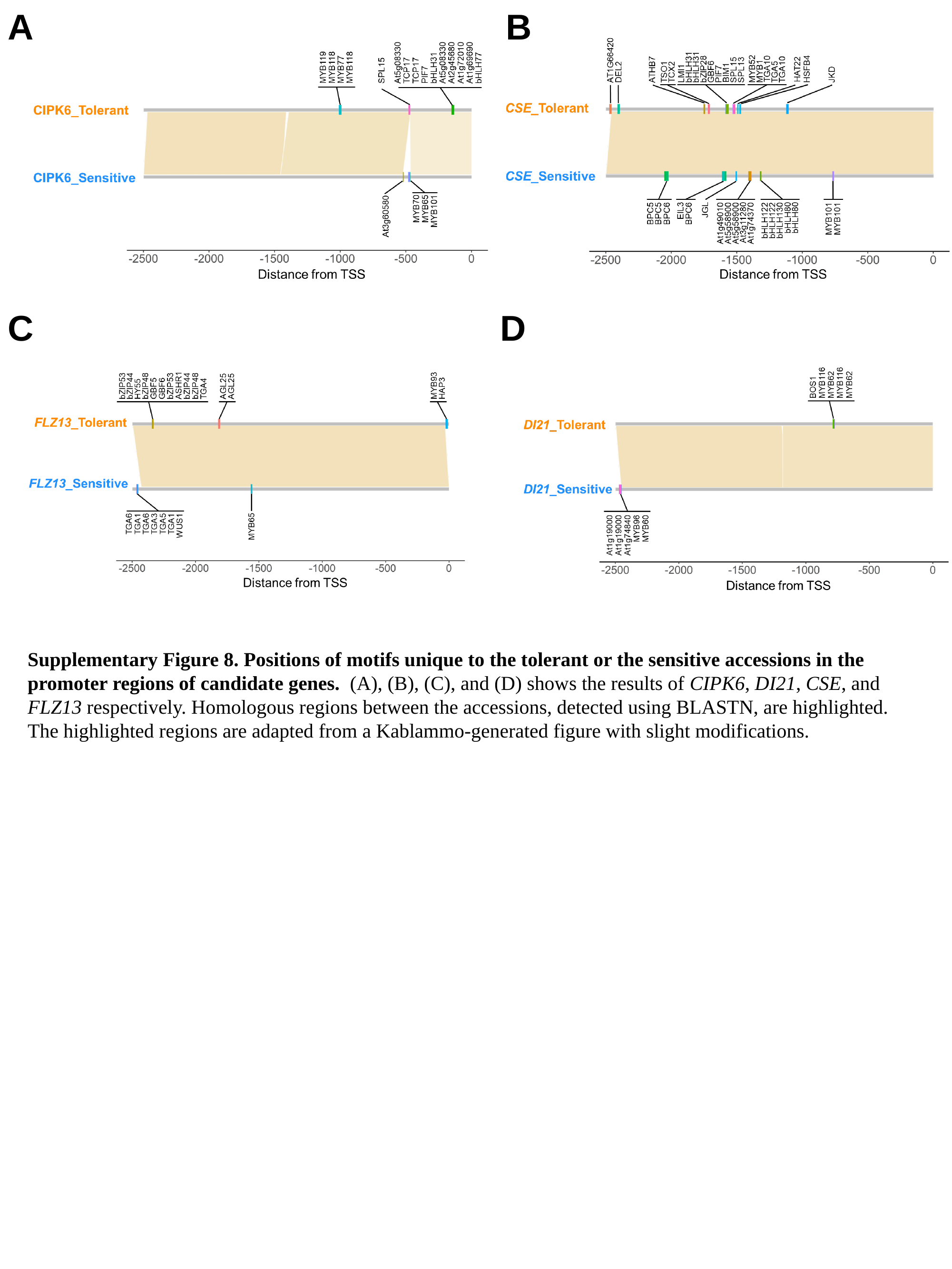

A
B
C
D
Supplementary Figure 8. Positions of motifs unique to the tolerant or the sensitive accessions in the promoter regions of candidate genes. (A), (B), (C), and (D) shows the results of CIPK6, DI21, CSE, and FLZ13 respectively. Homologous regions between the accessions, detected using BLASTN, are highlighted. The highlighted regions are adapted from a Kablammo-generated figure with slight modifications.

### Slide 9
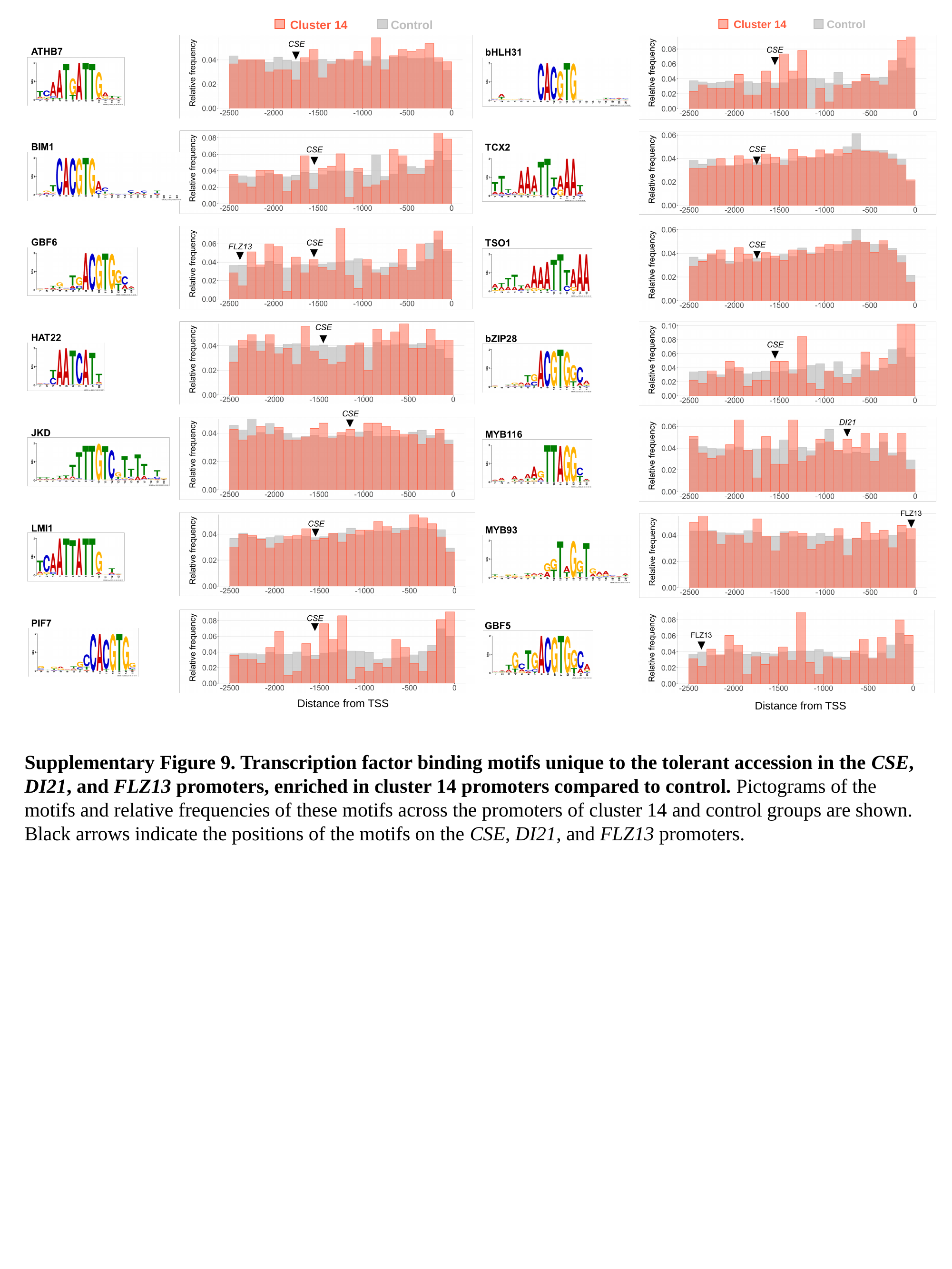

Cluster 14
Control
Cluster 14
Control
Distance from TSS
Distance from TSS
Supplementary Figure 9. Transcription factor binding motifs unique to the tolerant accession in the CSE, DI21, and FLZ13 promoters, enriched in cluster 14 promoters compared to control. Pictograms of the motifs and relative frequencies of these motifs across the promoters of cluster 14 and control groups are shown. Black arrows indicate the positions of the motifs on the CSE, DI21, and FLZ13 promoters.

### Slide 10
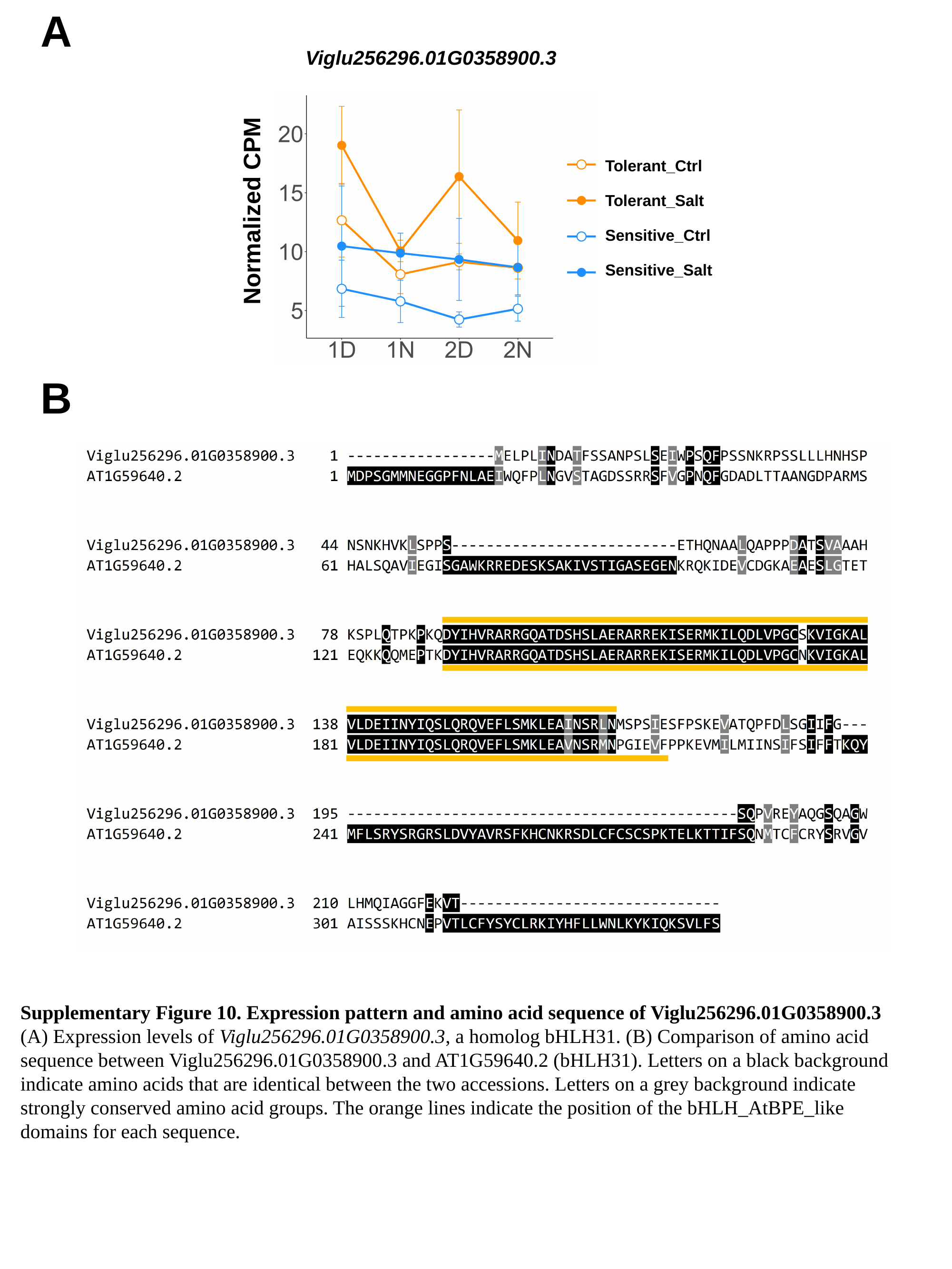

A
Viglu256296.01G0358900.3
Tolerant_Ctrl
Tolerant_Salt
Normalized CPM
Sensitive_Ctrl
Sensitive_Salt
B
Supplementary Figure 10. Expression pattern and amino acid sequence of Viglu256296.01G0358900.3
(A) Expression levels of Viglu256296.01G0358900.3, a homolog bHLH31. (B) Comparison of amino acid sequence between Viglu256296.01G0358900.3 and AT1G59640.2 (bHLH31). Letters on a black background indicate amino acids that are identical between the two accessions. Letters on a grey background indicate strongly conserved amino acid groups. The orange lines indicate the position of the bHLH_AtBPE_like domains for each sequence.
